# Naturally occurring cobalamin (B_12_) analogs can function as cofactors for human methylmalonyl-CoA mutase

**DOI:** 10.1101/2020.03.20.997551

**Authors:** Olga M. Sokolovskaya, Tanja Plessl, Henry Bailey, Sabrina Mackinnon, Matthias R. Baumgartner, Wyatt W. Yue, D. Sean Froese, Michiko E. Taga

## Abstract

Cobalamin, commonly known as vitamin B_12_, is an essential micronutrient for humans because of its role as an enzyme cofactor. Cobalamin is one of over a dozen structurally related compounds – cobamides – that are found in food and are produced by microorganisms in the human gut. Very little is known about how different cobamides affect B_12_-dependent metabolism in human cells. Here, we test *in vitro* how diverse cobamide cofactors affect the function of methylmalonyl-CoA mutase (MMUT), one of two cobalamin-dependent enzymes in humans. We find that, although cobalamin is the most effective cofactor for MMUT, multiple cobamides support MMUT function with differences in binding affinity (*K*_d_), binding kinetics (*k*_on_), and concentration dependence during catalysis (*K*_M, app_). Additionally, we find that six disease-associated MMUT variants that cause cobalamin-responsive impairments in enzymatic activity also respond to other cobamides, with the extent of catalytic rescue dependent on the identity of the cobamide. Our studies challenge the exclusive focus on cobalamin in the context of human physiology, indicate that diverse cobamides can support the function of a human enzyme, and suggest future directions that will improve our understanding of the roles of different cobamides in human biology.

## Introduction

Vitamins are diet-derived micronutrients that are essential for human health. Cobalamin (vitamin B_12_) is among a subset of vitamins that function as enzyme cofactors. Humans require cobalamin as a cofactor for two enzymes: methionine synthase (MS) and methylmalonyl-CoA mutase (MMUT, MCM in bacteria) (Figure 1A) (1). MS catalyzes the methylation of homocysteine, a reaction that is important not only because it produces methionine, a proteinogenic amino acid and precursor to the cofactor *S*- adenosylmethionine, but also for generating forms of tetrahydrofolate that are required for DNA synthesis (2). MMUT is a mitochondrial enzyme that catalyzes the reversible isomerization of (*R*)-methylmalonyl-CoA to succinyl-CoA, which is part of the propionate catabolism pathway in humans and is required for breaking down branched amino acids, odd-chain fatty acids, and the side-chain of cholesterol into the citric acid (TCA) cycle (3). Impairments in MMUT or MS activity, which can result from cobalamin deficiency in the diet, decreased cobalamin absorption (e.g. pernicious anemia), or inherited mutations in genes encoding MMUT, MS, and cobalamin trafficking proteins, lead to illnesses ranging from mild anemia to severe neurological dysfunction (4-8).

**Figure 1:**
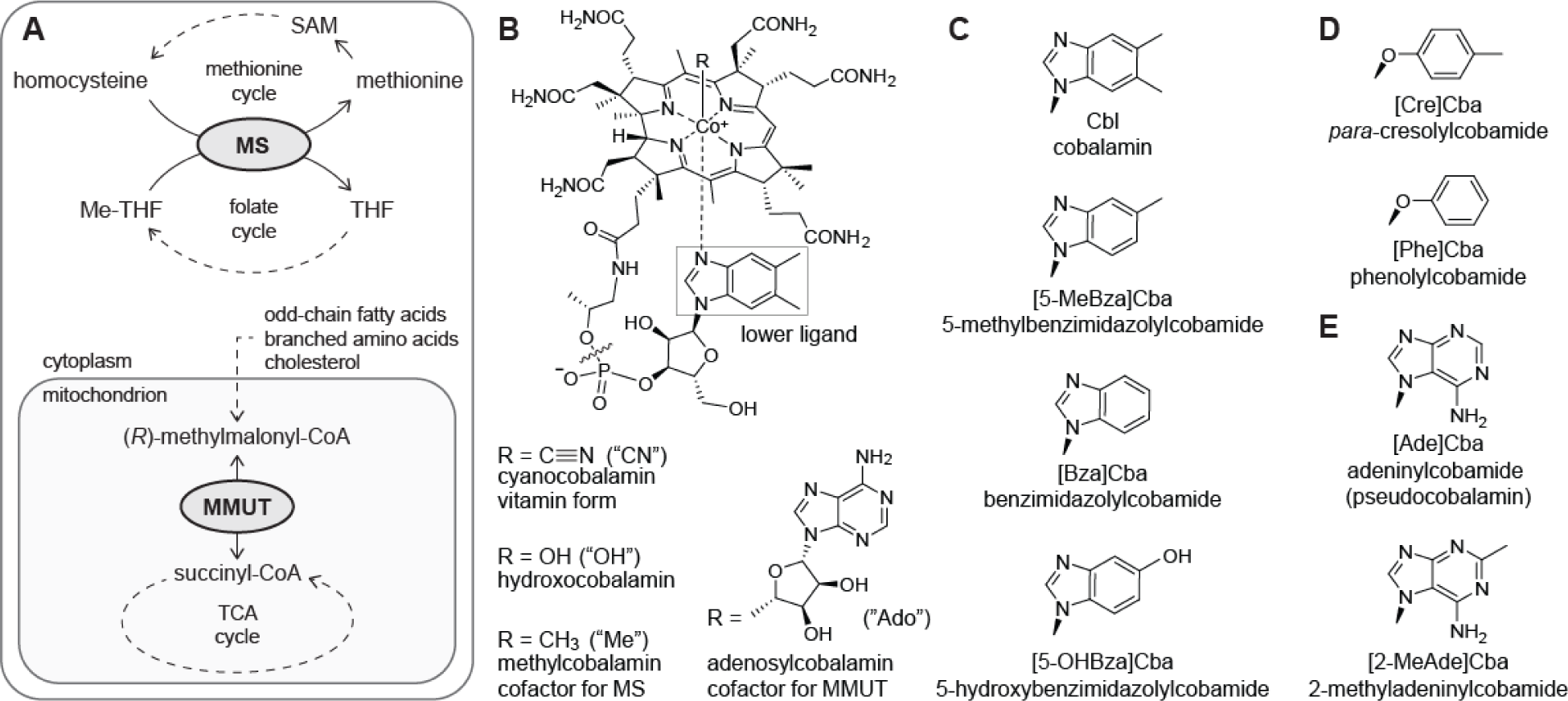
Cobalamin in human metabolism. (A) Diagram of metabolic pathways involving cobalamin in human cells. Dotted arrows indicate multiple reactions. MS (methionine synthase) and MMUT (methylmalonyl-CoA mutase) are the only cobalamin-dependent enzymes in humans. SAM, *S*- adenosylmethionine; (Me-)THF, (methyl-)tetrahydrofolate. (B) The structure of cobalamin. The upper ligand, R, varies for different enzymes; abbreviations listed in parentheses are used in the text when naming cobamides to specify the upper ligand. A wavy line delineates the part of cobalamin (including the lower ligand) that is absent in the precursor cobinamide. The lower ligand, boxed, is different in other cobamides. (C) Benzimidazolyl, (D) phenolyl, and (E) purinyl lower ligands found in cobamides.

Although it is found in animal tissues, cobalamin is produced exclusively by prokaryotes (9). Rather than synthesizing cobalamin, many bacteria and archaea produce cobalamin analogs that have the same core structure as cobalamin (Figure 1B) but differ in the identity of the nucleotide base, commonly referred to as the lower ligand (Figure 1B, boxed), which can be a benzimidazole, phenolic, or purine (Figure 1C-E) (10-15). Cobalamin and its analogs are collectively called cobamides (and are also known as corrinoid cofactors). In some environments, including the human gut, purinyl or phenolyl cobamides can be significantly more abundant than cobalamin itself (11,16). All cobamides share the same catalytic features, which include an upper ligand (*R* in Figure 1B) that varies for different chemical reactions; for the MMUT-catalyzed reaction, the upper ligand is 5’-deoxyadenosine (as in adenosylcobalamin, AdoCbl), while for MS it is a methyl group (as in methylcobalamin, MeCbl). Although it is not directly involved in catalysis, lower ligand structure affects the biochemistry of cobamide-dependent enzymes, including both MCM and MS (17-24). However, the differential effects of cobamides have been primarily studied in bacterial cobamide-dependent enzymes, and only to a limited extent in mammalian MS and MCM.

It is generally assumed that humans are unable to use cobamides other than cobalamin, in part because of the intricacy and selectivity of the human cobalamin uptake and trafficking system. Human intrinsic factor (IF) is a glycoprotein that captures various forms of cobalamin (including AdoCbl and cyanocobalamin, CNCbl, the vitamin form of cobalamin) in the intestine with up to femtomolar affinity and mediates uptake into ileal cells. IF has been reported to bind the purinyl cobamides adenosyladeninylcobamide (Ado[Ade]Cba, also known as pseudocobalamin) and adenosyl-2- methyladeninylcobamide (Ado[2-MeAde]Cba, also known as Factor A) six orders of magnitude more weakly than AdoCbl (25,26), and to have low affinity for cyano-*para*-cresolylcobamide (CN[Cre]Cba) and the cobamide precursor cobinamide (Figure 1B) (27). Human transcobalamin (TC), which subsequently binds cobalamin forms that have entered the bloodstream and facilitates uptake into various tissues, is also highly selective against cobinamide, but less selective than IF for AdoCbl over other cobamides (25-27). The selectivity of both proteins is considered important for preventing cobinamide, inactive cobamides, and cobamide degradation products from reaching MS and MMUT (18,28-32). Haptocorrin, a cobalamin-binding protein that binds structurally diverse cobamides and cobinamide with high affinity, might play an additional role in clearing non-cobalamin analogs from the body (25).

Despite the apparent selectivity of human cobamide uptake proteins, cobalamin analogs have been detected in patient serum samples (33,34) and in human liver (35). One possible source of cobalamin analogs, including other cobamides, in humans is the diet. Some shellfish, and, most notoriously, edible cyanobacteria such as *Spirulina*, have high content of cobamides other than cobalamin (36), and bacterial species associated with the production of fermented foods such as yogurt synthesize alternate cobamides (37). Additionally, some human gut bacteria that reside in the small intestine, the region of the gut where cobalamin is absorbed, synthesize alternate cobamides (38) (cobamides produced by bacteria in the large intestine are not spatially bioavailable to humans (39,40)). Although IF is selective for cobalamin, it binds cyano-5-methylbenzimidazolylcobamide (CN[5-MeBza]Cba) with similar affinity as CNCbl, and IF affinity for cyanobenzimidazolylcobamide (CN[Bza]Cba) and cyano-5-hydroxybenzimidazolylcobamide (CN[5-OHBza]Cba) is reported to be no more than 2-fold lower than its affinity for CNCbl (26,27). Thus, these benzimidazolyl cobamides are likely able to enter ileal cells. Moreover, despite its low affinity for purinyl cobamides, IF may bind these analogs to some extent at physiological concentrations (25), especially if they are enriched in the diet. Furthermore, TC is less selective than IF among benzimidazolyl cobamides (26,27), and binds purinyl cobamides with higher affinity than IF (25), so any cobamides that do enter the bloodstream may be bound by TC. The consequent possibility of different cobamides reaching human cells warrants investigation of the effects of these cobamides on human cobalamin-dependent enzymes. While MS has been shown to be active with multiple cobamides *in vitro* (31), whether or not cobamides with diverse lower ligands are suitable cofactors for MMUT has not, to our knowledge, been reported.

Here, we investigate the ability of human MMUT to use cobamide cofactors other than cobalamin *in vitro*. We find that MMUT binds to and is active with several cobamides from different structural classes, with differences in binding affinity, binding rate, and activity kinetics. We additionally characterize the ability of a set of MMUT missense variants to use different cobamides (41). These MMUT variants represent a subset of hundreds of mutations in MMUT that are found in patients with the inherited metabolic disorder methylmalonic aciduria (MMA) (41,42). Using a collection of 14 natural and unnatural cobamides and cobinamide, we screen the activity of six MMUT variants to look for improved activity *in vitro*. Although we find that the activity of MMUT variants with different cobamides remains well below that of the wildtype enzyme, this investigation demonstrates that both wildtype and MMUT variants are able to use several cobamides as cofactors, contrary to the assumption that cobalamin is the only suitable cobamide for humans.

## Results

### MMUT binds many cobamides with varying affinity

To determine whether MMUT is able to use cobamides other than cobalamin, we heterologously expressed MMUT in *E. coli* and assayed the ability of the purified enzyme to bind cobamides from different structural classes. We measured quenching of intrinsic protein fluorescence to determine equilibrium dissociation constants (*K*_d_ values) for eight cobamides and the cobamide precursor cobinamide, which lacks a lower ligand (Figure 1B). The *K*_d_ calculated for AdoCbl, the native cofactor of MMUT, by this method was 0.08 ± 0.03 µM (Figure 2A, C). Previous studies of MMUT report AdoCbl *K*_d_ values of 0.04 µM (43) and 0.27 ± 0.11 µM (44), and bacterial MCM orthologs are reported to bind AdoCbl with affinities within this range (24,32), lending credence to our *K*_d_ determination. Using the same binding assay, we found that MMUT binds several benzimidazolyl and phenolyl cobamides in addition to AdoCbl (Figure 2A, B), although structural differences within these classes resulted in differences in binding affinity spanning up to two orders of magnitude (Figure 2C, e.g. the *K*_d_ values of AdoCbl and Ado[Bza]Cba, which differ by two methyl groups, differ by ∼25-fold). Surprisingly, MMUT appeared to have higher affinity for Ado[Cre]Cba than AdoCbl. In contrast, MMUT did not bind purinyl cobamides to a significant extent at micromolar concentrations (Figure 2B, C). Consistent with previous reports focused on bacterial MCM orthologs (24,32), MMUT bound Ado-cobinamide, but with lower affinity than AdoCbl (Figure 2A, C). The small overall fluorescence decrease observed upon Ado-cobinamide binding could be due to the conformation of the enzyme-cofactor complex in the absence of a lower ligand, which likely differs relative to other cobamides; the locations of aromatic residues that are most likely to report on binding are shown in Figure S1A.

**Figure 2:**
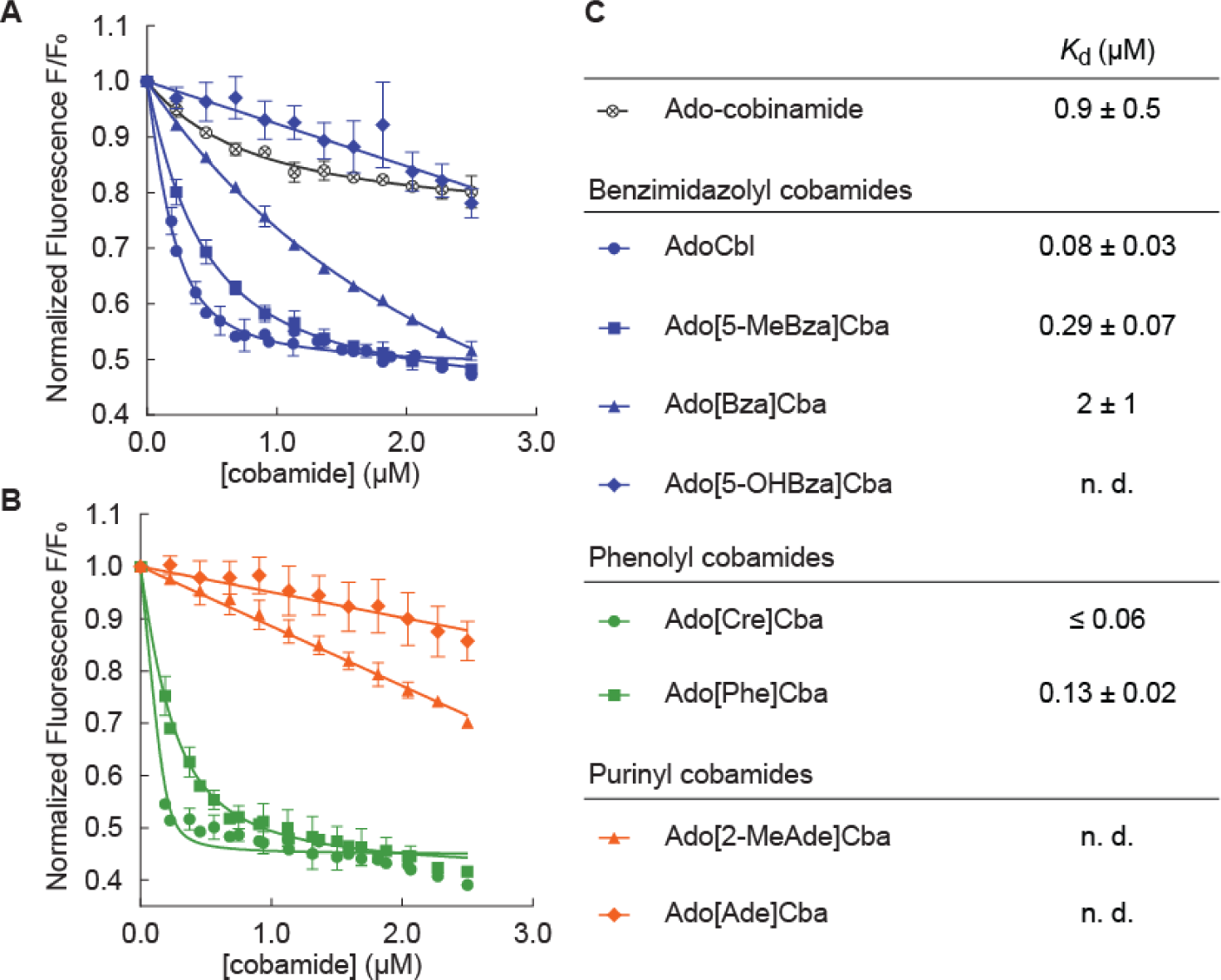
Binding of structurally diverse cobamides to MMUT. Fluorescence decrease of MMUT (0.2 µM) reconstituted with (A) benzimidazolyl cobamides (blue) and cobinamide (gray), and (B) phenolyl (green) and purinyl (orange) cobamides, at the following concentrations: 0.19 – 2.5 µM for AdoCbl and phenolyl cobamides, and 0.23 – 2.5 µM for all other cofactors. Data points represent the mean and standard deviation of three technical replicates from a single experiment. (C) *K*_d_ values are presented as the average and standard deviation of five or more technical replicates across at least two independent experiments. An upper limit of the *K*_d_ is reported for Ado[Cre]Cba, which bound too strongly to measure a *K*_d_ value in the given experimental conditions. “n. d.,” not determined, indicates that binding was too weak to determine *K*_d_.

We focused our attention on the cobamides with the highest affinities for MMUT to examine whether lower ligand structure affects cofactor binding kinetics. Using stopped-flow fluorescence spectroscopy, we determined binding rates (*k*_on_) for AdoCbl and the two phenolyl cobamides. Interestingly, we discovered that the *k*_on_ of Ado[Cre]Cba was about six times higher than the *k*_on_ of AdoCbl and adenosylphenolylcobamide (Ado[Phe]Cba), which had similar binding rates (Figure 3). The fast *k*_on_ of Ado[Cre]Cba may explain its high binding affinity for MMUT compared to AdoCbl and Ado[Phe]Cba.

**Figure 3:**
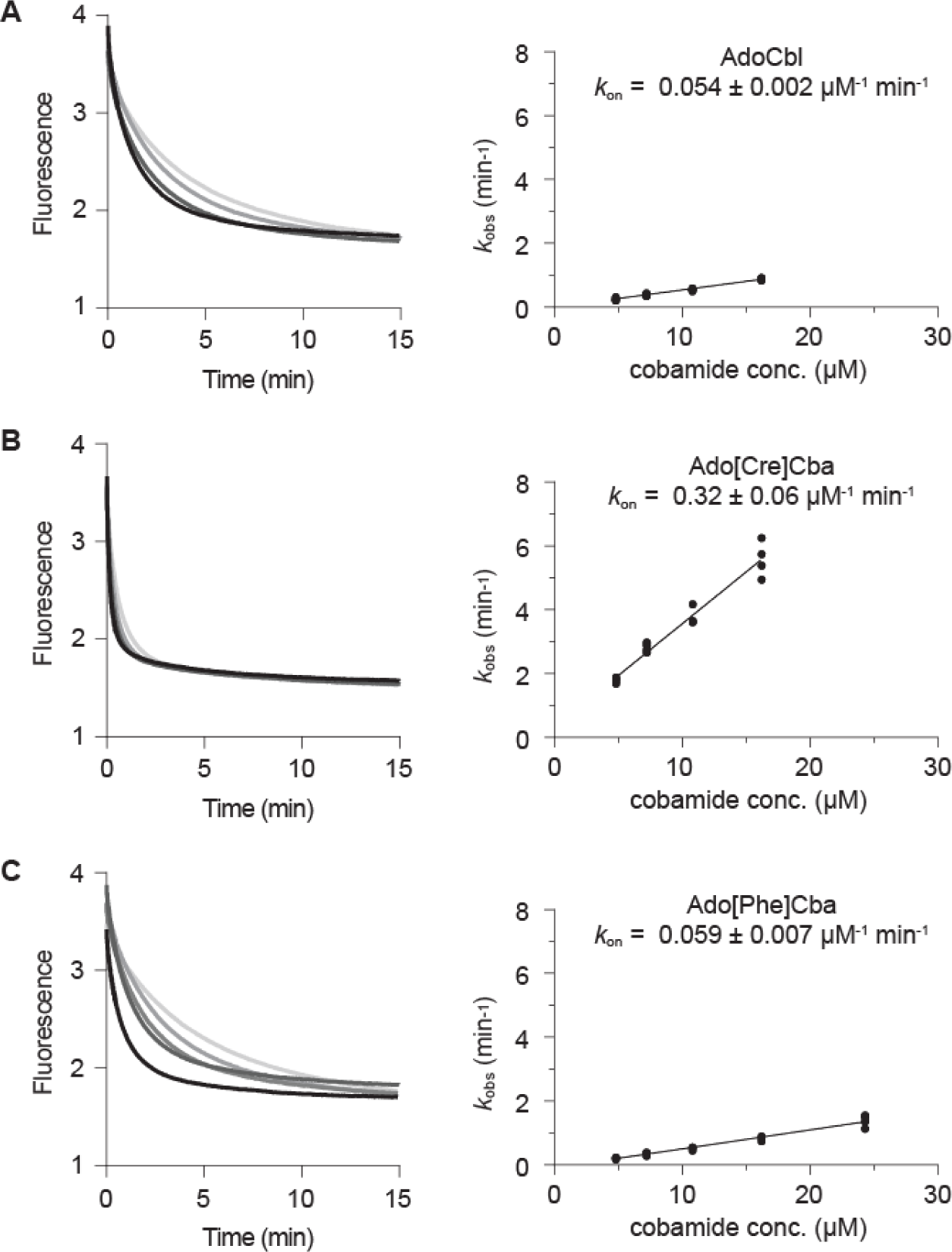
Binding kinetics of AdoCbl and phenolyl cobamides. On the left, time course of the fluorescence decrease of MMUT (0.2 µM) upon addition of (A) AdoCbl, 4.8 – 16.2 µM; (B) Ado[Cre]Cba, 4.8 – 16.2 µM; and (C) Ado[Phe]Cba, 4.8 – 24.3 µM (concentrations increase from light to dark). Data were fitted to an exponential decay function to determine binding rates (*k*_obs_), which were plotted as a function of cobamide concentration (on the right) to calculate *k*_on_. *k*_on_ values are the average and standard deviation of four technical replicates.

### Multiple cobamides support MMUT activity, with differences in apparent K_*M*_

Given that several cobamides bind MMUT, we next considered the possibility that MMUT activity can be supported by cobamides other than AdoCbl *in vitro*. We characterized the activity of recombinant MMUT reconstituted with cobamides for which the enzyme had high affinity, including both benzimidazolyl and phenolyl cobamides. Using a coupled spectrophotometric assay we found that MMUT was active with all of the cobamides we tested: AdoCbl, Ado[5-MeBza]Cba, Ado[Bza]Cba, Ado[Cre]Cba and Ado[Phe]Cba. Interestingly, however, the concentration required to achieve half-maximal activity, *K*_M, app_, of the cobamides differed drastically, spanning two orders of magnitude (Figure 4). AdoCbl had the lowest *K*_M, app_ (0.04 ± 0.02 µM, in agreement with a previously reported value of 0.050 µM (45)), and the other benzimidazolyl cobamides had *K*_M, app_ values within 10-fold of AdoCbl (Figure 4A, C). The 10-fold difference in *K*_M, app_ between AdoCbl and Ado[Bza]Cba is consistent with a previous report for MCM purified from sheep kidney (18). While the *K*_M, app_ of Ado[Cre]Cba was in the range of the benzimidazolyl cobamides, Ado[Phe]Cba had a *K*_M, app_ 100-fold higher than AdoCbl (Figure 4B, C). The high *K*_M, app_ values of both phenolyl cobamides relative to AdoCbl were unexpected, given their high binding affinities for MMUT.

**Figure 4:**
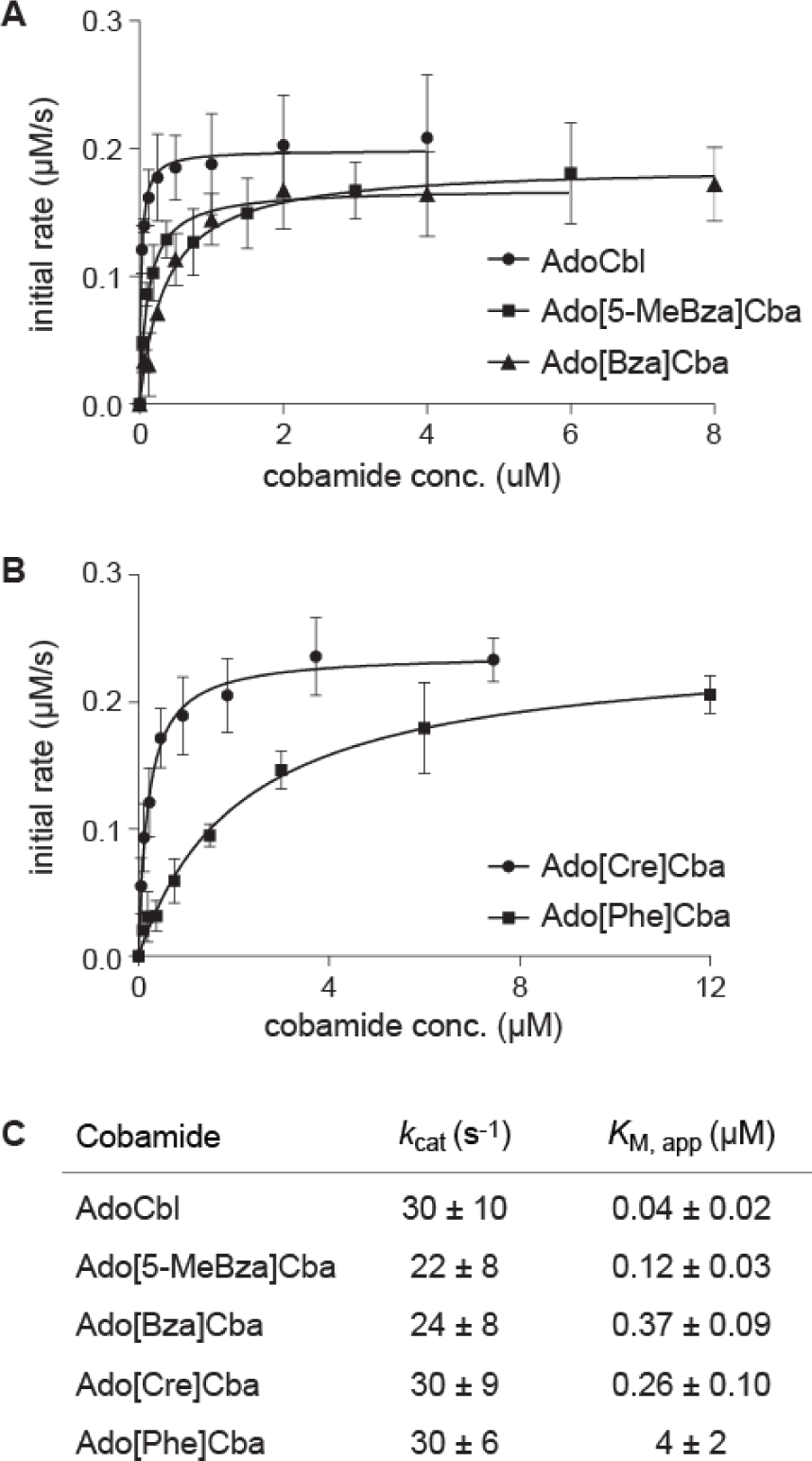
MMUT kinetics. Isomerization of methylmalonyl-CoA (4 mM) to succinyl-CoA by MMUT (10 nM) was measured after MMUT reconstitution with (A) benzimidazolyl and (B) phenolyl cobamides (AdoCbl: 0.031 – 4 µM, Ado[5-MeBza]Cba: 0.047 – 6 µM, Ado[Bza]Cba: 0.063 - 8 µM, Ado[Cre]Cba: 0.058 – 7.5 µM, Ado[Phe]Cba: 0.094 – 12 µM). Data points and error bars represent the mean and standard deviation, respectively, of three technical replicates from one experiment. *K*_M, app_ and *k*_cat_ values are reported in (C) as the average and standard deviation of five or more replicates from at least two independent experiments and two biological samples.

### Disease-associated MMUT variants with defects in AdoCbl K_*M*_ also have impaired activity with other cobamides

Increased *K*_M, app_ of AdoCbl is a biochemical defect associated with some disease-causing variants of MMUT (41). Since we discovered that four cobamides besides AdoCbl support wildtype MMUT activity but have distinct binding and kinetic properties, we considered the possibility that cobamides other than cobalamin might also support activity of MMUT mutant variants and potentially suffer less of a *K*_M_ defect than AdoCbl. If this were the case, such cobamides could be further investigated as a potential disease therapy for MMA. We therefore screened a library of cobamides and cobinamide for the ability to enhance the activity of six MMA-causing MMUT variants associated with *K*_M_ defects (41). Three of these variants have amino acid substitutions located in the B_12_-binding domain of the enzyme (G648D, V633G, G717V), while the other substitutions (P86L, Y100C, Y231N) are located in the substrate-binding domain near the cofactor or near the MMUT dimer interface (41,45) (Figure S1B). We determined the specific activity of wildtype (WT) MMUT and each MMUT variant reconstituted with nine naturally occurring cobamides (Figure 5A, compounds A-H and M), cobinamide, and six “unnatural” cobamide analogs that we previously biosynthesized for structure-function studies (24) (Figure 5A, compounds J-L and N-P).

**Figure 5:**
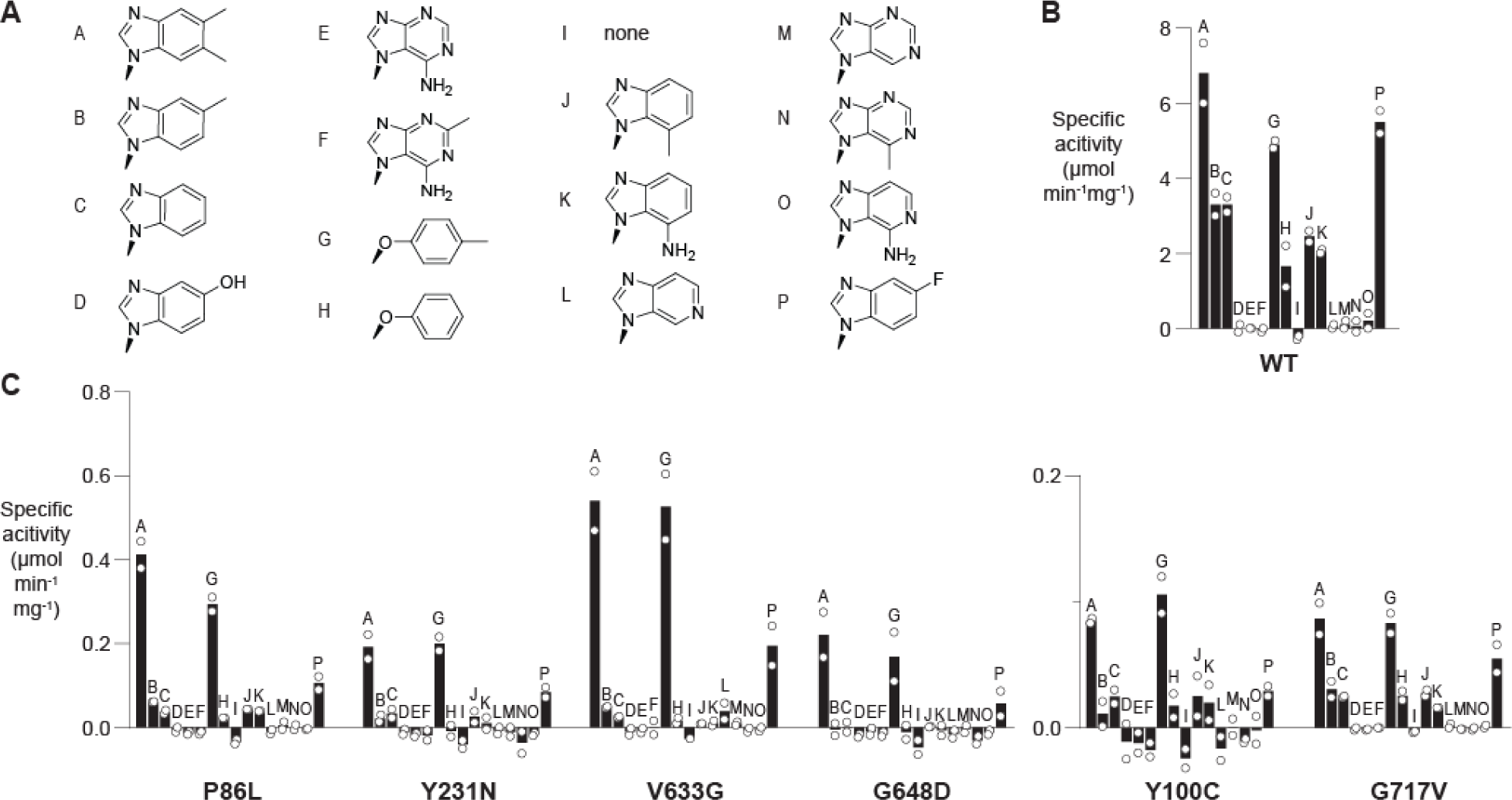
Activity screen of MMUT variants associated with disease. (A) Lower ligands of cobamides screened in this experiment, including cobamides introduced in Figure 1 (A-I in this figure) as well as J: 7-methylbenzimidazolylcobamide, K: 7-aminobenzimidazolylcobamide, L: 6- azabenzimidazolylcobamide, M: purinylcobamide, N: 6-methyladeninylcobamide, O: 3- deazaadeninylcobamide, and P: 5-fluorobenzimidazolylcobamide. Specific activity of (B) MMUT WT (0.01 µM) and (C) mutant variants (0.1 µM, except G717V at 1 µM) reconstituted with different adenosylated cobamides and cobinamide (all at 1 µM) was determined after 30 minutes of activity. Each letter (A – P) corresponds to a compound assigned in (A). Bars represent the average specific activity across two independent experiments (circles). Note the differences in the y-axis scales between WT and mutated proteins.

Consistent with the results of our kinetic assay (Figure 4), WT MMUT (Figure 5B) was active with AdoCbl, Ado[5-MeBza]Cba, and Ado[Bza]Cba (compounds A, B, C), as well as phenolyl cobamides (compounds G, H), and was additionally active with all three of the unnatural benzimidazolyl cobamide analogs (compounds J, K, P). No activity was observed with Ado[5-OHBza]Cba (compound D) or purinyl cobamides (compounds E, F, M, N), likely due to their low binding affinities (Figure 2), or with azabenzimidazolyl cobamides (compounds L, O).

As expected, all of the variants had significantly reduced specific activity with AdoCbl compared to WT MMUT (Figure 5C). None of the cobamides tested rescued MMUT variant activity to levels comparable to the WT enzyme. However, Ado[Cre]Cba (compound G) supported activity to a large extent in all of the variants, and some mutants also had appreciable activity with adenosyl-5- fluorobenzimidazolylcobamide (compound P) (Figure 5C). Notably, most of the variants, with the exception of G717V, partially or entirely lost the ability to use benzimidazolyl cobamides other than AdoCbl (compounds B, C, J, K) relative to the WT enzyme. This implies that if other benzimidazolyl cobamides reach human cells in patients with some of these mutations, the deleterious effects of the mutations could potentially be exacerbated.

## Discussion

Two human enzymes, MS and MMUT, require the cofactor cobalamin. The metabolic functions of MS and MMUT are essential, and for this reason cobalamin deficiency can be fatal (4,5). Humans have evolved a complex, high-affinity uptake, processing, and delivery system to capture cobalamin in food and escort it to MS and MMUT. In addition to cobalamin, however, the human diet contains other cobamides, as well as cobamide precursors and degradation products (28,36,37,46). The selectivity of the cobamide uptake system is important for protecting human cells from analogs that inhibit MS and MMUT. However, it is conceivable that cobamide cofactors other than cobalamin reach human cells, given the ability of IF and TC to bind multiple cobamides (26,27). Supporting this possibility are reports of cobalamin analogs in human tissues (33-35) and the findings that orally and subcutaneously administered cobamide analogs can be found in mammalian tissues (26,30). It is therefore important to understand how alternate cobamides impact the biochemistry of cobamide-dependent enzymes in humans. Prior to this study, whether cobamides with diverse lower ligands are suitable cofactors for MMUT had not been investigated.

Here, we report that MMUT is able to use several, but not all, cobamides as cofactors *in vitro*. Variations in lower ligand structure have multiple effects on MMUT function. Lower ligand structure strongly impacts cobamide binding to MMUT, which is consistent with previous reports of selective cobamide binding by bacterial MCM orthologs (24). We found that benzimidazolyl and phenolyl cobamides bound MMUT with affinities spanning over two orders of magnitude, and purinyl cobamides did not bind MMUT at micromolar concentrations. All cobamides that bound MMUT supported the catalytic activity of the enzyme, but had up to 100-fold differences in *K*_M, app_. Our initial expectation was that *K*_M,app_ values of cobamides would equal their *K*_d_ values; since cobamides remain bound to MMUT for multiple reaction cycles during which the cofactor is continuously regenerated (47,48), half-maximal activity is expected when MMUT is half-saturated with cobamide (i.e., at *K*_d_). The differences between *K*_M, app_ and *K*_d_ that we observed may therefore be indicative of distinct effects of lower ligands on the catalytic cycle of MMUT, in addition to their effect on binding.

We also measured binding kinetics of different cobamides to MMUT and found that lower ligand structure affects the binding rate of cobamide cofactors. The high *k*_on_ of Ado[Cre]Cba compared to AdoCbl is reminiscent of a previous report that the reaction of 2-methyleneglutarate mutase, a bacterial cobamide-dependent enzyme that catalyzes a similar rearrangement to the one catalyzed by MMUT, has a shorter lag time following addition of Ado[Cre]Cba compared to AdoCbl (21). Fast binding of Ado[Cre]Cba to MMUT and 2-methyleneglutarate mutase could be explained by the absence of an intramolecular coordinate bond in this cobamide; coordination between the lower ligand and the cobalt ion, which is present in AdoCbl in solution (Figure 1B), must be disrupted for the cofactor to bind both enzymes (49). This may also explain why Ado-cobinamide, which lacks a lower ligand entirely, binds a bacterial MCM ortholog more rapidly than AdoCbl (32). The fact that Ado[Phe]Cba also has a slower binding rate to MMUT despite lacking a coordinated lower ligand, however, suggests that the structure of the lower ligand itself plays a role in binding kinetics.

In the X-ray crystal structure of MMUT bound to AdoCbl, the hydrophobic lower ligand of AdoCbl is located within a highly hydrophobic binding pocket of the enzyme (50) (Figure S1C). Impaired binding of Ado[5-OHBza]Cba and purinyl cobamides to MMUT may therefore be explained by the absence of stabilizing hydrophobic interactions between MMUT and the more polar lower ligands of these cobamides. We previously observed a similar pattern in MCM from the bacterium *Sinorhizobium meliloti* (*Sm*MCM), and identified nitrogen atoms within the six-membered ring of purinyl lower ligands as the structural feature that impaired purinyl cobamide binding (24). These ring nitrogens may similarly interfere with binding of purinyl cobamides to MMUT; the absence of enzyme activity after MMUT reconstitution with any cobamides containing nitrogens in the six-membered ring (compounds L-O, in addition to E and F, in Figure 5) is consistent with this speculation. Comparing both studies reveals that human MMUT and *Sm*MCM, which have 61% sequence identity, are remarkably similar in their relative affinities for different cobamides, and are unlike other bacterial MCM orthologs from *Escherichia coli* and *Veillonella parvula* (24). The similarities between MMUT, a mitochondrial enzyme, and *Sm*MCM could reflect their evolutionary relationship, as mitochondria are thought to share a more recent common ancestor with *S. meliloti* (an α-proteobacterium) than with *E. coli* (a γ-proteobacterium) and *V. parvula* (phylum Firmicutes) (51); however, more experimental evidence would be required to support this hypothesis.

Given their functional differences, administration of diverse cobamides to humans could be considered as a possible therapy if MMUT variants associated with disease have improved activity with cobamides other than cobalamin. Among the six MMUT variants that we tested, none of the 14 cobamides in our collection rescued activity to a significant extent compared to cobalamin, although at least two other cobamides supported the activity of each mutant. While addition of high concentrations of cobalamin has been shown to improve the activity of MMUT variants classified as *mut*^-^ (41), some mutations in MMUT cannot be rescued by cobalamin addition (*mut*^0^) and are associated with more severe pathologies and death; however, whether any other cobamides support the activity of *mut*^0^ variants has not been tested. Thus, there is room for further investigating cobamides as therapies for inherited disorders of cobalamin.

Therapeutic use of alternate cobamides would depend on the ability of these cobamides to reach the MMUT active site in a living organism, which requires that they be absorbed into the bloodstream, transported to various tissues (these initial steps can potentially be bypassed by direct injection of free cobamides or cobamides pre-bound to TC, which has been done in rabbits (26)), and recognized by multiple cobalamin trafficking proteins in human cells (4,52). For example, in cells cobalamin is loaded to MMUT directly by an adenosyltransferase enzyme, which activates the cofactor by installing the 5’- deoxyadenosyl upper ligand, and this transfer is gated by a GTPase that facilitates release of inactivated cofactor (53-55). The extent to which these enzymes are sensitive to lower ligand structural variation is currently unknown. The need to investigate lower ligand selectivity in cobamide trafficking and activation is highlighted by apparent discrepancies between *in vitro* characterization of purified mammalian cobamide-dependent enzymes and experiments administering cobamides to live animals. Specifically, Kolhouse *et al*. demonstrated that while human MS was active with multiple cobalamin analogs *in vitro* (31), subcutaneous administration of most of these analogs in rats did not support MS activity *in vivo* (30). Similarly, while our data demonstrate that MMUT is active with Ado[Bza]Cba, subcutaneous administration of OH[Bza]Cba in rats appeared to mildly inhibit MMUT activity (30). The discrepancy between *in vitro* results with purified enzymes and experiments in live animals (with the caveat that MS and MMUT activity assays were performed on the human enzymes, while the animal studies were performed in rats) may be attributable to the many proteins that interact with cobalamin prior to its binding MS or MMUT. Therefore, fully understanding the impact of diverse cobamides on human physiology, and assessing their therapeutic relevance, requires further expanding our knowledge of the biochemical impacts of lower ligand structure on absorption, trafficking, and activation of these cofactors.

## Experimental procedures

### Protein expression and purification

Two preparations of MMUT were used in this work. (1) MMUT was expressed with an N-terminal hexahistidine (6xHis) tag in *E. coli* BL21(DE3)pLysS from the pMCM-2 expression plasmid kindly provided by María Elena Flores (47). The expression strain was grown at 37 °C to an optical density at 600 nm of 0.6, cooled on ice for 20 min, and protein was expressed for 19 h at 16 °C after induction with 1 mM isopropyl-*β*-D-thiogalactopyranoside (IPTG). Cells were lysed by sonication in 50 mM sodium phosphate pH 8.0, 100 mM NaCl, with 0.5 mM PMSF, 1 µg/mL leupeptin, 1 µg/mL pepstatin, and 1 mg/mL lysozyme. Protein was purified using nickel affinity chromatography and dialyzed into 25 mM Tris-HCl pH 7.5, 300 mM NaCl, and 10% glycerol. The final concentration of 6xHis-tagged MMUT was determined by absorbance at 280 nm using the extinction coefficient 72,310 M^-1^ cm^-1^. (2) Wildtype and mutant MMUT variants were expressed with C-terminal 6xHis tags and purified as originally described in Froese *et al*. (50) with modifications provided in Forny *et al*. (41).

MMUT prepared by both methods (1) and (2) was used for equilibrium binding assays. The binding affinity of MMUT for AdoCbl was indistinguishable between preparations using both methods. All binding kinetics and activity assays were conducted using MMUT prepared by method (2).

### Cobamide production

All cobamides besides cobalamin were produced in bacterial cultures and purified as previously described (13,24,56,57). Cobamides and cobinamide were chemically adenosylated to obtain active (5’- deoxyadenosylated) forms (24,58,59), which was the form of the cofactor used for all experiments reported in this manuscript.

### Cobamide binding assays

Cobamide equilibrium binding affinities were determined by measuring quenching of intrinsic protein fluorescence caused by cofactor binding, as previously described (24,32). Briefly, MMUT (0.2 µM) was combined with varying concentrations of cobamides (between 0.19 and 2.5 µM; see Figure 2) in a 96-well plate, and fluorescence was measured on a Tecan Infinite M1000 Pro plate reader using excitation/emission wavelengths of 282 nm/340 nm. Because ligand concentrations were not in significant excess of MMUT, data were fit to a quadratic binding equation that accounts for the depletion of free ligand during binding (24,60).

Binding kinetics were measured by stopped-flow fluorescence spectroscopy using the Kintek AutoSF-120 stopped flow fluorimeter. Fluorescence emission was detected with a 320 nm long pass filter (320FG01, Andover Corporation) at an excitation wavelength of 282 nm (± 0.12 nm). Temperature and buffer conditions were identical to the equilibrium binding assay. The final concentration of MMUT after mixing was 0.2 µM, and cobamide concentrations added were 4.8, 7.2, 10.8, and 16.2 µM (and 24.3 µM for Ado[Phe]Cba). To determine binding rates (*k*_obs_), fluorescence decreases were fitted to a first-order exponential decay equation, following subtraction of MMUT fluorescence decrease upon addition of buffer. *k*_on_ was determined by plotting *k*_obs_ as a function of cobamide concentration (where *k*_on_ is the slope).

### MUT activity assays

Enzyme kinetics were measured by coupling MMUT activity to thiokinase (which hydrolyzes succinyl-CoA to succinate and CoA), and spectrophotometric detection of CoA using dithionitrobenzoate (DTNB), as previously described (24,61). We verified that the concentrations of thiokinase and coupling reagents used in the assay were not rate limiting in previous work (24). The assay was modified as follows: the concentration of (*R*)-methylmalonyl-CoA was fixed at 4 mM, MMUT at 10 nM, and cobamide concentrations were varied across reactions as specified in Figure 4. (*R*)-Methylmalonyl-CoA was enzymatically synthesized (24).

A modified version of the coupled activity assay was used to screen activity of mutated MMUT variants. The enzymes were pre-incubated with cobamides, at 1.25X final concentration, in 100 mM Tris, 50 mM phosphate pH 7.5, on ice, for 30 minutes (32) in a 384-well plate. The plate was transferred to 30 °C and a 5X mixture containing thiokinase, MgCl_2_, ADP, and methylmalonyl-CoA was added to initiate the reaction. Final concentrations of reagents are as previously described (24), with the following adjustments: cobamide concentration, 1 µM (a saturating concentration of AdoCbl for WT activity); (*R*)- methylmalonyl-CoA, 2 mM; MMUT WT, 0.01 µM; MMUT G717V 1 µM; all other MMUT variants, 0.1 µM. DTNB was omitted from the reaction mixture, as it was found to inhibit protein activity on timescales longer than those used to measure initial rates. After 30 minutes, a sample of the reaction mixture was removed and immediately combined with DTNB (2.5 mM). Absorbance at 412 nm was measured on a BioTek Synergy 2 plate reader and concentration of CoA was calculated using the extinction coefficient 14,150 M^-1^ cm^-1^.

## Acknowledgements

We thank members of the Taga lab and Judith Klinman for helpful discussions, Eric Greene for help with stopped-flow experiments, Kathryn Quanstrom and Kenny Mok for their contributions during the early stages of this project, María Elena Flores for the MMUT expression plasmid, and Andreas Martin and Krishna Niyogi for sharing their equipment. This work was supported by National Institutes of Health grants R01GM114535 and DP2AI117984 to M.E.T and Swiss National Sciences Foundation [31003A_175779] to M.R.B. Structural Genomics Consortium is a registered charity (Number 1097737) that receives funds from AbbVie, Bayer Pharma AG, Boehringer Ingelheim, Canada Foundation for Innovation, Eshelman Institute for Innovation, Genome Canada, Innovative Medicines Initiative (EU/EFPIA) [ULTRA-DD grant no. 115766], Janssen, Merck & Co., Novartis Pharma AG, Ontario Ministry of Economic Development and Innovation, Pfizer, São Paulo Research Foundation-FAPESP, Takeda, and Wellcome Trust [092809/Z/10/Z].

---

**Figure S1:**
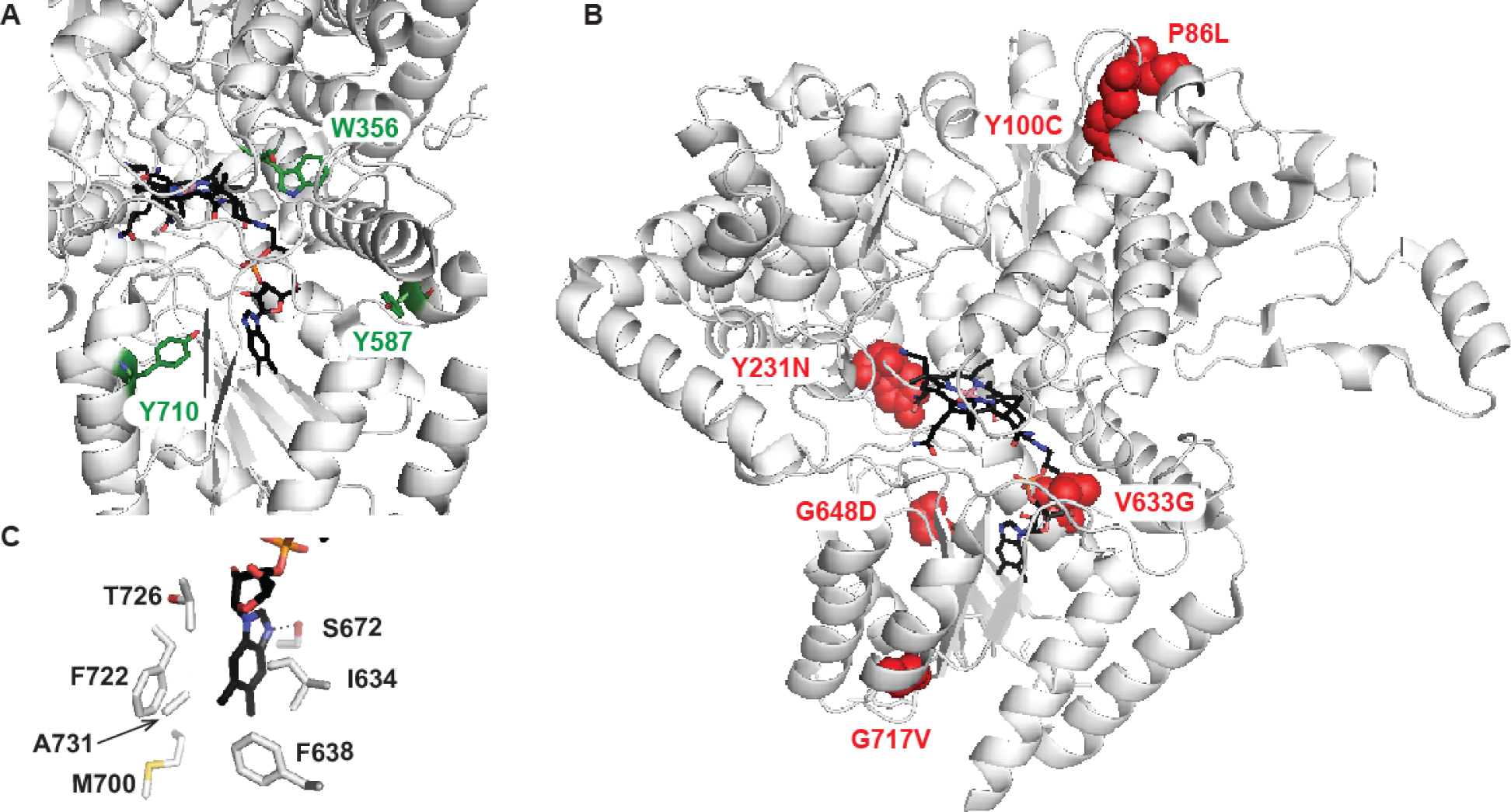
X-ray crystal structure of MMUT bound to cobalamin (PDB: 2XIJ) (50). In white is the enzyme; in black, cobalamin. (A) In green, residues hypothesized to contribute to fluorescence quenching upon cofactor binding. (B) In red, locations of disease-associated mutations assayed in Figure 5. (C) Residues immediately surrounding the lower ligand of cobalamin.

